# Two- and three-dimensional tracking of *MFA2* mRNA molecules in mating yeast

**DOI:** 10.1101/2020.07.02.185355

**Authors:** Polina Geva, Konstantin Komoshvili, Stella Liberman-Aronov

## Abstract

Intracellular mRNA transport contributes to the spatio-temporal regulation of mRNA function and localized translation. In the budding yeast, *Saccharomyces cerevisiae*, asymmetric mRNA transport localizes ∼30 specific mRNAs including those encoding polarity and secretion factors, to the bud tip. The underlying process involves RNA binding proteins (RBPs), molecular motors, processing bodies (PBs), and the actin cytoskeleton. Recently, pheromone a-factor expression in mating yeast was discovered to depend upon proper localization of its mRNA, *MFA2*. *MFA2* mRNAs in conjunction with PBs cluster at the shmoo tip to form “mating bodies”, from which a-factor is locally expressed. The mechanism ensuring the correct targeting of mRNA to the shmoo tip is poorly understood.

Here we analyzed the kinetics and trajectories of *MFA2* mRNA transport in living, alpha-factor treated yeast. Two-(2D) and three-dimensional (3D) analyses allowed us to reconstruct the granule tracks and estimate granule velocities. Tracking analysis of single *MFA2* mRNA granules, labeled using a fluorescent aptamer system, demonstrated three types movement: vibrational, oscillatory and translocational. The mRNA granule transport was complex; a granule could change its movement behavior and composition during its journey to the shmoo. Processing body assembly and the actin-based motor, Myo4p, were involved in movement of *MFA2* mRNA to the shmoo, but neither was required, indicating that multiple mechanisms for translocation were at play. Our visualization studies present a dynamic view of the localization mechanism in shmoo-bearing cells.

## Introduction

Intracellular mRNA transport is one of the most important post-transcriptional mechanisms of gene regulation. The proper transport of mRNA plays an important role in localized translation, which is critical for establishing cell polarity and subcellular function. In eukaryotes, for example in fibroblasts, neurons and drosophila oocytes, it is necessary for many biological processes, including polarized growth, division, development and differentiation [1,2].

There are at least three mechanisms for localizing mRNAs in the cytoplasm: (a) directed transport to a place, (b) local protection from degradation, and (c) diffusion and entrapment by a localized anchor [3,4]. In the first mechanism, mRNAs are packaged into ribonucleoprotein (RNP) particles that are directly transported and anchored at the site of localization [5,6]. Transport of these RNP particles depends on specific *cis*-acting element(s), motor proteins and cytoskeletal filaments [7]. In the second mechanism, the RNAs are protected from degradation at one site and are highly susceptible to degradation in other parts of the cell. In the third mechanism, mRNAs freely diffuse throughout the cell but upon reaching the site of localization, are unable to leave, producing a concentration “hot spot” [8].

In budding yeast, *Saccharomyces cerevisiae*, ∼30 specific mRNAs localize at the bud tip, including those for polarity and secretion factors, [9–11]. Formation of transport-competent RNPs is initiated via the recognition of *cis*-regulatory elements present in RNA molecules by specific RNA-binding proteins (RBPs), termed trans-acting factors [4,12]. The regulatory elements are usually located within the 3’UTR [9,13]. The directed transport of *ASH1, SRO7* and *IST2* mRNA in budding yeast is mediated by RNP complexes. After transcription, maturated RNPs are exported from the nucleus, whereupon their movement through the cytoplasm to their destinations occurs on the cytoskeleton by molecular motor proteins [14] and reviewed in [3]. Localization and transport of *bicoid* mRNA in *Drosophila* and *Xdazl* mRNA in *Xenopus* oocytes as well as that of *β*-actin mRNA in fibroblast and neuronal cells have varied mechanisms principally based on RNP assembly and cytoskeletal support [4].

Understanding the dynamic behavior of mRNA movement may shed light on the mechanism by which these mRNAs localize. Methods for the dynamic study of native mRNA in living cells now exist. Current, technical developments in intracellular RNA imaging allow for observation of the movement of single mRNA molecules in living cells in real time [7,15,16]. One popular approach is to insert a linker site for RBP within the 3’UTR of the mRNA of interest. The RBP is conjugated to a fluorophore that enables microscopic tracking of individual mRNA transcripts in living cells [7,17]. In order to track single mRNA movement in real time, it is important to achieve high sensitivity for single molecule detection and fast image acquisition. A sufficient tracking range is crucial to identify the type of motion. The total number of frames in the image sequence determines the statistical accuracy of the analysis.

Most studies were performed in 2D, but it is highly desirable to extend the technology into 3D since most biological processes occur within a volume [18]. Recently, single mRNA particles in live budding yeasts were tracked in three dimensions using a microscope with a double-helix point spread function [6,19]. Simultaneous observation of *ASH1* and *IST2* mRNA revealed directed co-transport from the mother cell into the bud, followed by corralled movements of both mRNAs once they reached the bud cortex [19].

In the cytoplasm, mRNAs are always chaperoned by RBPs that create RNP complexes [20]. These complexes can be quite variable in size, properties and function, which is dependent on its contents. Processing bodies (PBs) are a special type cytoplasmic RNP complex, primarily composed of translationally repressed mRNAs and proteins related to mRNA decay, suggesting roles in post-transcriptional regulation [21,22]. PBs may also play a role in mRNA transport as evidenced by the various types of PB movements in different cell types. [23–25]. The mRNAs transported within PBs to specific locations are protected from degradation and preliminary translation by RBPs present in the complex which are also regulated in their transport to specific destinations [25,26].

In budding yeast, mRNP transport is associated with multiple Myo4p motors via the activity of the RBPs, She2p and adaptor She3p [3,27]. Transported mRNPs accompany cortical endoplasmic reticulum (ER) using the SHE family proteins [29,30]. RNP granules and ER movement is consistent with the speed generated by a motor. The *Ash1* mRNA is contained in RNPs that move at velocities that vary between 0.2–0.44 µm/s [7], which are consistent with a myosin V motor [31]. Cell-specific RNA velocity estimates provide a natural basis for intracellular mRNA behavior during cell cycle. The PBs containing mRNA/RBPs complexes promote not only transport from mother to daughter cell, but also mRNA local translation required for gene function [27,32].

Recently, attention has turned to mRNA transport in mating yeast. The advantage of this system over budding yeast, is that in order to mate, haploid cells of the opposite genotype (MATα/a) create an asymmetrically growing long protrusion called a shmoo in response to pheromone treatment. It allows for the investigatation of mRNA transport behavior and its trajectory over long distances, similar to axonal transport in neuronal cells [33–35]. Our previous research uncovered the role of mRNA transport and localization in the regulation of aF translation in mating yeast. We demonstrated that in response to pheromone treatment, cytoplasmic *MFA2* mRNA is transported to the shmoo tip where it is locally translated [27]. Only a small fraction of the mRNA granules located in mother cells shows direct rapid movement from the cell body to the shmoo tip where it accumulates as one large granule named “mating body.” These transported mRNPs are associated with the PB marker, Dcp2. *MFA2* mRNA is eventually released for local translation of a-factor (aF). aF is exported out of the cells by the ATP binding cassette (ABC) transporter Ste6p at the shmoo tip for activation of effective mating with a potential partner [32]. For comparison, an unrelated mRNA, *PGK1* (Phosphoglycerate Kinase 1) that is translated to a glycolytic enzyme, distributes randomly throughout the cell. No directed *PGK1* mRNA transport to the shmoo was found using two-dimensional video analysis [27]. The mechanisms responsible for correctly targeting proteins to the shmoo tip are not well understood.

The *MFA2* mRNA showed distinct subgroups of the granules, which were different in location in the cell, size, and motion properties. In our previous studies, we carried out two-dimensional imaging studies (2D); here, we investigated the dynamics of all types of *MFA2* mRNA granules with a high spatial resolution during shmoo growth using four dimensional studies (4D), including z-stack and time intervals, which were transformed to 3D for analysis that should give more accurate tracking information about position, behavior and velocity of the granules than 2D. We addressed several questions: What trajectories do they exhibit? Three-dimensional measurements are essential to extract full information about mRNA dynamics during movement including physical properties. What factors participate in their movement? What role does a motor Myo4p play in *MFA2* mRNA delivery to the shmoo? To address these questions, we described and classified mRNA movements in *S*. *cerevisiae* after stimulating MATa cells expressing fluorescently labeled *MFA2* mRNA with α-factor mating pheromone.

## Results

We investigated the dynamics of *MFA2* mRNA granules labeled with U1A-GFP in α-factor treated, WT yeast cells (ySA056, Table 5). In parallel, we examined the behavior of *PGK1* mRNA, that expressed transcript from a housekeeping gene, *PGK1*, (ySA061, Table 5) as a control unrelated to mating. We used a previously developed system consisting of two plasmids encoding *MFA2*-U1A binding sites and U1A-GFP [17,36]. To allow for live-cell imaging of individual mRNA granules, each transcript insert contained 16 repeated aptamer fragments in its 3’UTR region to enable recognition by U1A RNA binding protein fused with GFP. The aptamer-tagged mRNAs were localized using fluorescence microscopy. All cells under observation formed only a single shmoo, suggesting that all responsive cells possessed unidirectional polarity. Randomly chosen cells with different stages of shmoo development were used for imaging. Within a given cell, individual granules moved independently and in different directions, indicating that their motion was not due to bulk movement of the cytoplasm.

### Range of movement of *MFA2* and *PGK1* mRNA in 3D

To understand the mechanism of localizing *MFA2* granules to the shmoo tip, we compared the movement of *MFA2* and *PGK1* granules in 3D. Similar to what we have published previously in 2D [27], there were three essential types of *MFA2* granule motion in 3D by using Image Analysis Software (Imaris) (Fig. 1, Methods, S1_ Movie.mov): vibrational, oscillatory and translocational that have distinct sizes, locations in the cells, and behavior. Only two types of *MFA2* mRNA granules were motile: oscillatory and translocational. The vibrational and oscillatory *MFA2* mRNA granules were mostly observed in the main cell body, whereas translocated mRNA granules were more motile and were delivered to shmoo. Figure 1A shows a representative image of several *MFA2* mRNA containing granules, that vary in size from 0.8 to 0.2 μm. Size was dependent on the number of copies of mRNA molecules per granule and/or RBP types associated with it.

**Fig. 1.**
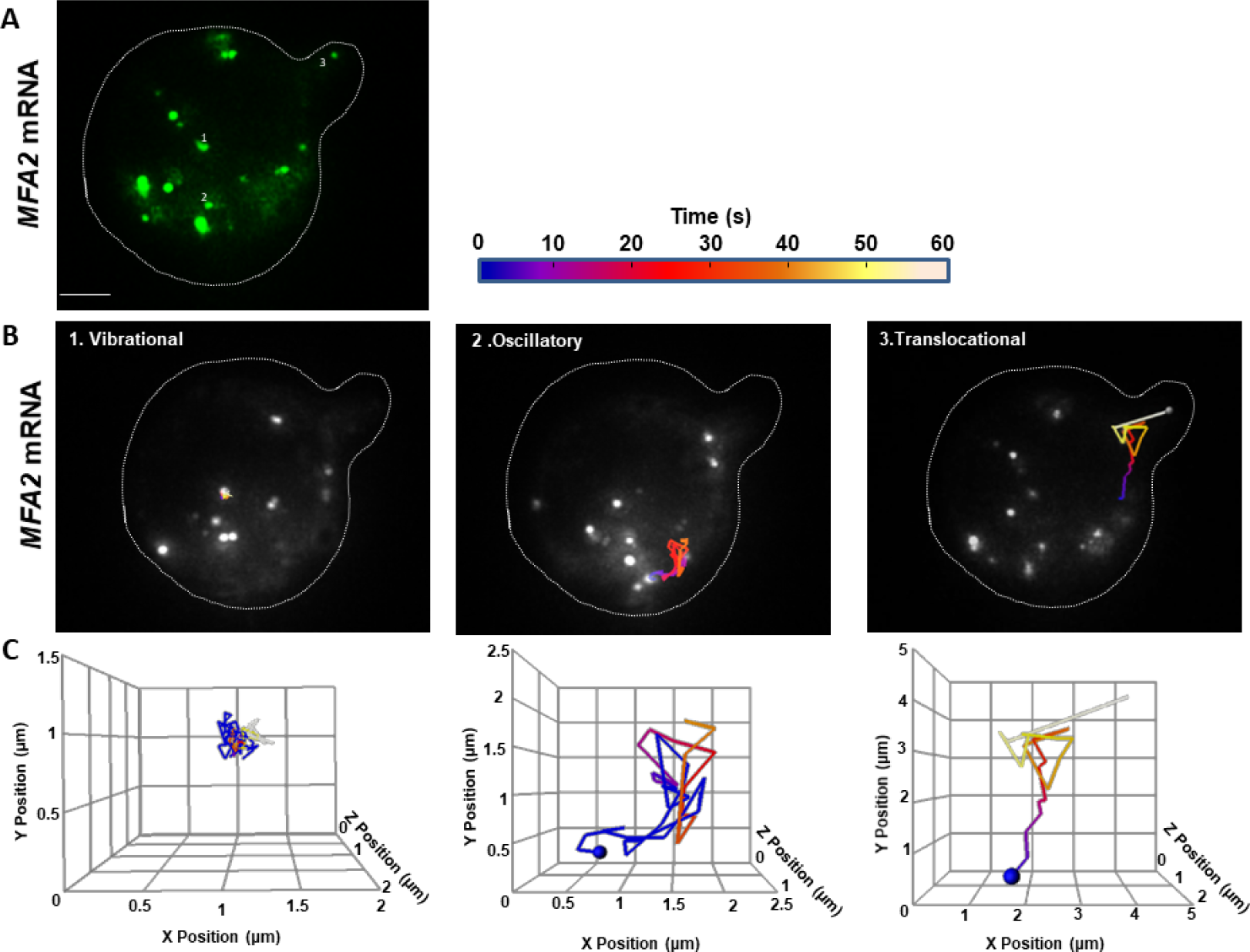
*MFA2* granules exhibited three types of motion in 3D tracking. **(A)** U1A-GFP labeled *MFA2* mRNA in a yeast cell were visualized after treatment with α-factor for 2 hours. Broken line outlines the cell border with a shmoo developing on the upper right side. Numbers refer to particles tracked in (B)-(C). Scale bar = 2 µm also applies to (B). **(B)** Tracks of three particles display distinct behavior movements: vibrational, oscillatory, and translocational were superimposed on images of the cell at various fixed times. Tracks were constructed from images recorded every 4 s, consolidating 6 z-stacks with a depth of 0.5 µm for each cell. **(C)** Tracks of the *MFA2* granules shown in (B) are shown replotted in 3D. The axes were arbitrarily resized to illustrate the movement space of the granules. Blue spheres mark the initial positions of the granules. Track duration was 60s.

A “track displacement” parameter, defined as the change in position of an mRNA granule relative to its starting point, was used to classify movement type of *MFA2* RNP (Table 1). For normalization, this parameter and the overall distance traveled (track length), were calculated for 60s periods (see Methods). For *MFA2* mRNA granules undergoing translocational movement, track displacements ranged from 1.5 to 4.0 μm, with a mean of 2.3 μm. Translocational movement appeared to be restricted to low intensity granules, 0.2 to 0.5 μm in diameter. The lowest intensity mRNA granules (0.2 μm) were actively transported to shmoo tips (Figure 1B and C) and they contained up to 10 mRNA transcripts, according to previous estimates [37]. Oscillating granules traveled the same overall distance (track length) in the same period as translocating granules, but their track displacement was twofold less (1.1 μm). Their movement was restricted to a local area inside of the cell body and they only moved a slight distance away from the starting point, as compared to the translocational granules. For granules undergoing vibrational movement, track displacement was limited to 0.8 µm and was fourfold less as compared to translocated granules. They appeared to be attached to a place and displayed characteristics of immotile granules. Granules undergoing oscillatory or vibrational movement ranged widely in size; our analysis focused on those varying from 0.2 to 0.8 μm in diameter.

**Table 1.** Range of motion parameters for *MFA2* and *PGK1* granules tracked in 3D

| Type of motility | Vibrational |  | Oscillatory |  | Translocational |  |
| --- | --- | --- | --- | --- | --- | --- |
| Type of mRNA | *MFA2* | *PGK1* | *MFA2* | *PGK1* | *MFA2* | *PGK1* |
| Track length ($\mu\text{m}$) | 2.5±1.4 | 5.5±2.6 | 6.5±3.1 | 14.6±3.9 | 6.4±2.3 | --- |
| Track displacement ($\mu\text{m}$) | 0.5±0.2 | 0.6±0.2 | 1.1±0.3 | 1.5±0.4 | 2.3±0.7 | --- |

A total of 150 granules were analyzed in 47 cells for 60s periods. Track lengths and track displacements are given as mean ± SEM.

Unlike the *MFA2* granules that moved a significant distance from the cell body to the shmoo (Fig. 1B-C, Table 1), *PGK1* granules were smaller, ranging in size from 0.1 – 0.5 μm. They were more motile, although they moved only within limited sub-regions and were never observed to translocate to other cell compartments over periods of 60 s (Fig. 2, S2_ Movie.mov). However, track lengths for oscillatory *PGK1* granules were 2.3-fold greater than for *MFA2* granules (P<0.001), due to the more frequent occurrence of local jumps (see following sections). Vibrational track displacements of *MFA2* and *PGK1* granules were similar, but track length may have been several times larger for the latter (Table 1). The distinct behavior of *PGK1* mRNA granules could be due to their having a different granule composition, which is known to be important for regulation of granule stability, translational rate and degradation.

**Fig. 2.**
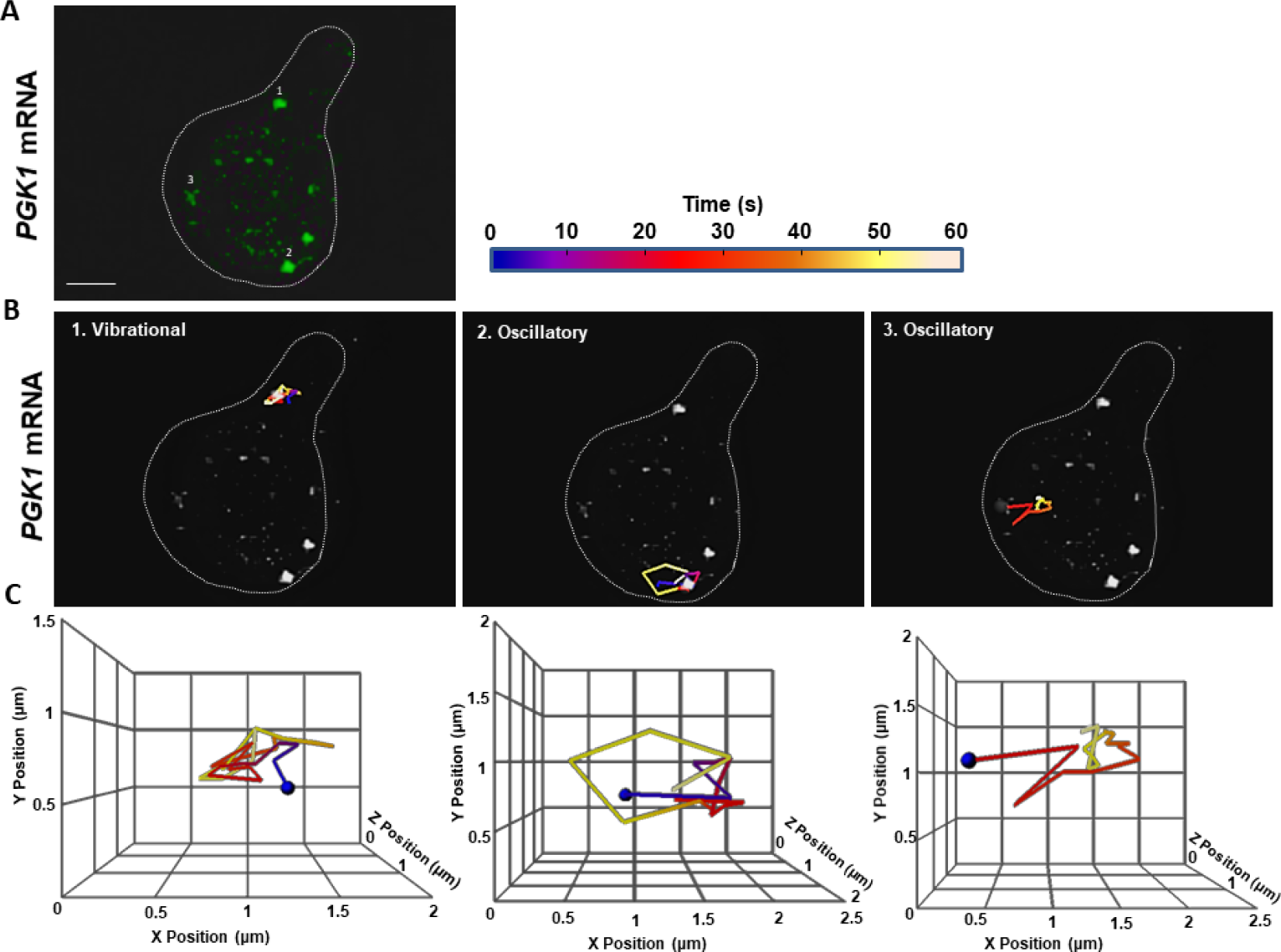
Only vibrational and oscillatory motions occurred in *PGK1* granules in 3D. The explanation is similar to that for Figure 1, except that this cell expressed U1A-GFP labeled *PGK1* mRNA. *PGK1* granules did not exhibit translocational movement. Scale bar = 2 µm.

To summarize, according to 3D analysis, most *PGK1* mRNA granules showed strong, local motile behavior, while none translocated to the shmoo. *MFA2* mRNA granules also showed restricted to local dynamic movement albeit to a somewhat lesser extent, but in addition, some low intensity *MFA2* granules underwent directed movement toward the shmoo.

### Comparison of *MFA2* granule movements in 2D and in 3D

To find out whether a significant amount of movement occurred on a fast time scale, mean velocity and track displacement rate of the fastest subset of translocating *MFA2* granules (0.2 -0.4 μm in diameter) were examined in 2D using a very sensitive camera mounted on a standard fluorescent microscope that allowed volume shooting of the granules without z stack scanning per frame (see Methods), (Fig. 3, S3_ Movie.mov). Sacrificing information about granule movement in the z-axis reduced frame capture time to 0.055-0.070 s. For translocating granules, the mean velocity calculated from the total track length in 2D and the track duration was 1.5 µm/s. In contrast, “track displacement rate” determined as 2D track displacement divided by track duration was only 0.04 μm/s (Table 2). The discrepancy between mean velocity and track displacement rate reflected the circuitous route traveled as well as several periods of aimless wandering.

**Table 2.** 2D vs 3D comparison of translocating *MFA2* granules

| Tracking mode | Frame period<br>(s) | Track displacement<br>rate ($\mu\text{m/s}$) | Mean velocity<br>($\mu\text{m/s}$) |
| --- | --- | --- | --- |
| 2D | 0.055-0.070 | 0.04±0.01 | 1.5±0.7 |
| 3D | 2-4 | 0.03±0.01 | 0.15±0.06 |

10 granules were analyzed for each tracking mode. Mean velocity was calculated as the total track length divided by the observation period and track displacement rate as the track displacement divided by the observation period. Observation periods varied from 15 to 250 s. Values are given as mean ± SEM.

**Fig. 3.**
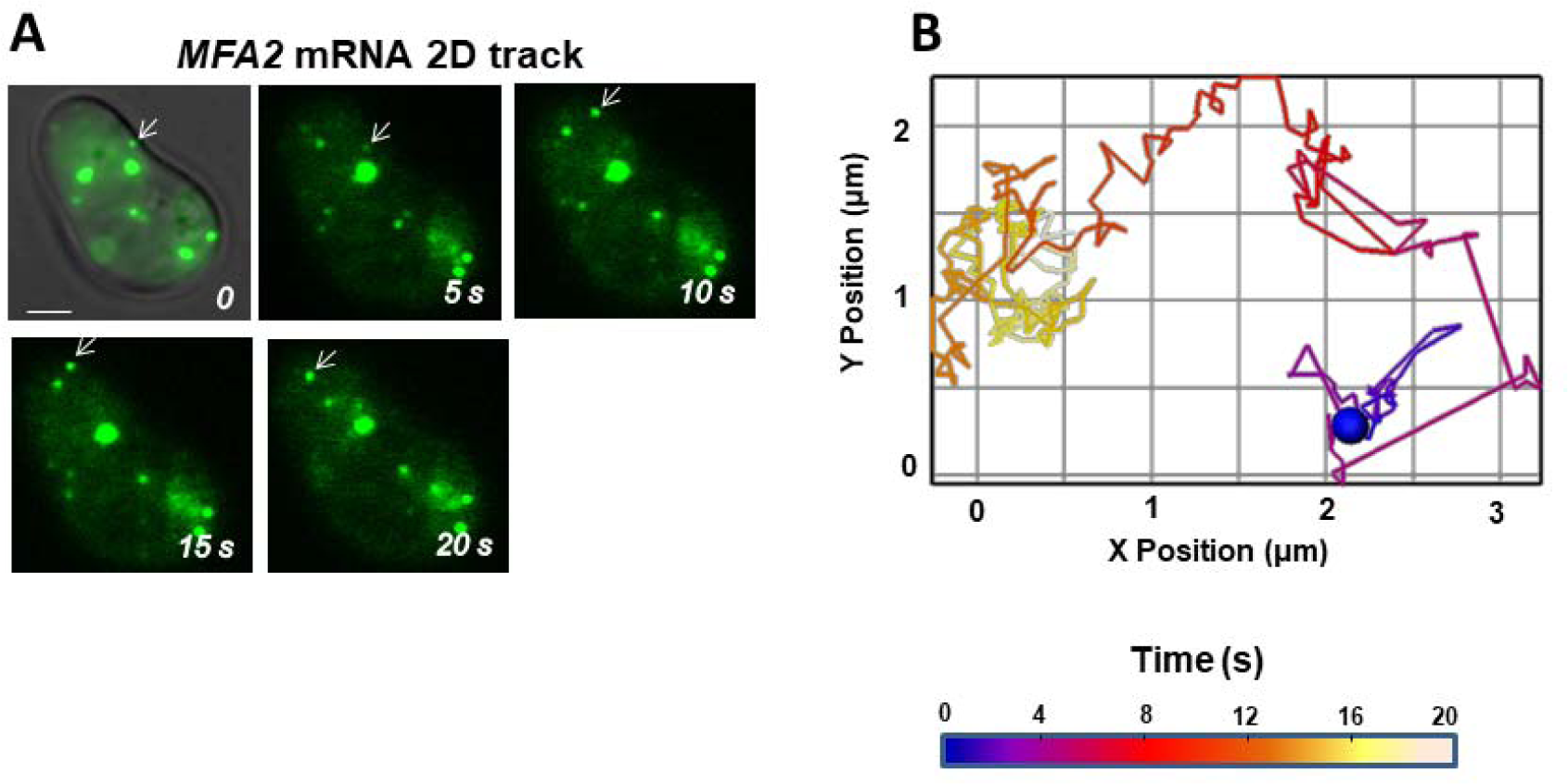
High time resolution tracking in 2D revealed extensive *MFA2* granule movement. **(A)** Cells expressing mRNA (green) treated with α-factor for 2 h were examined by fluorescence microscopy. There was a rudimentary shmoo on the upper left side of the cell. White arrow indicates the motile, low-intensity mRNA transport granule analyzed in (B). Scale bar = 2 µm. **(B)** Over most of its journey, the granule changed directions incessantly and took a very indirect route to the shmoo. The color scale refers to the time. Track displacement rate was 0.01 μm/s. Mean velocity was 0.73 μm/s.

Motile, low-intensity mRNA granules 0.2 -0.5 μm in diameter were analyzed for 3D mRNA transport in real-time using confocal microscopy (S4_ Movie.mov). The cells typically had a thickness of 3-4 μm. These granules were similar to the granules analyzed in 2D but included some that were somewhat larger and/or slightly more intense. The lowest intensity, fastest-moving *MFA2* granules that also had directed movement as shown previously in 2D here and in [27] were not accessible for the 3D analyses. Z scanning increased significantly the frame acquisition time to 2-4 s and we were able to follow more accurately, the details of the RNP tracks as compared to 2D. Figure 4A and S4<u>_</u> Movie.mov show an example of 3D tracking at 2 s/frame, of an mRNA granule as it moved from the neck to the tip of the shmoo. The average velocity for 10 granules was 0.15 μm/s and the average of the track displacement rate was 0.03 μm/s (Table 2). Compared to the analysis in 2D, the speed estimated in 3D was 10-fold slower, while the track displacement rate was similar. To rule out a systematic bias in granule selection by size and fluorescence intensity in 2D vs 3D imaging, the 2D velocities of four translocational granules were recalculated for 2 s intervals, yielding a range of 0.08-0.2 µm/s that corresponded more closely to values obtained with 3D analysis. Thus, the discrepancy in mean velocity reflected a limitation in the rate of frame capture, which was 60 to 80 times faster in 2D than in 3D imaging.

**Fig. 4.**
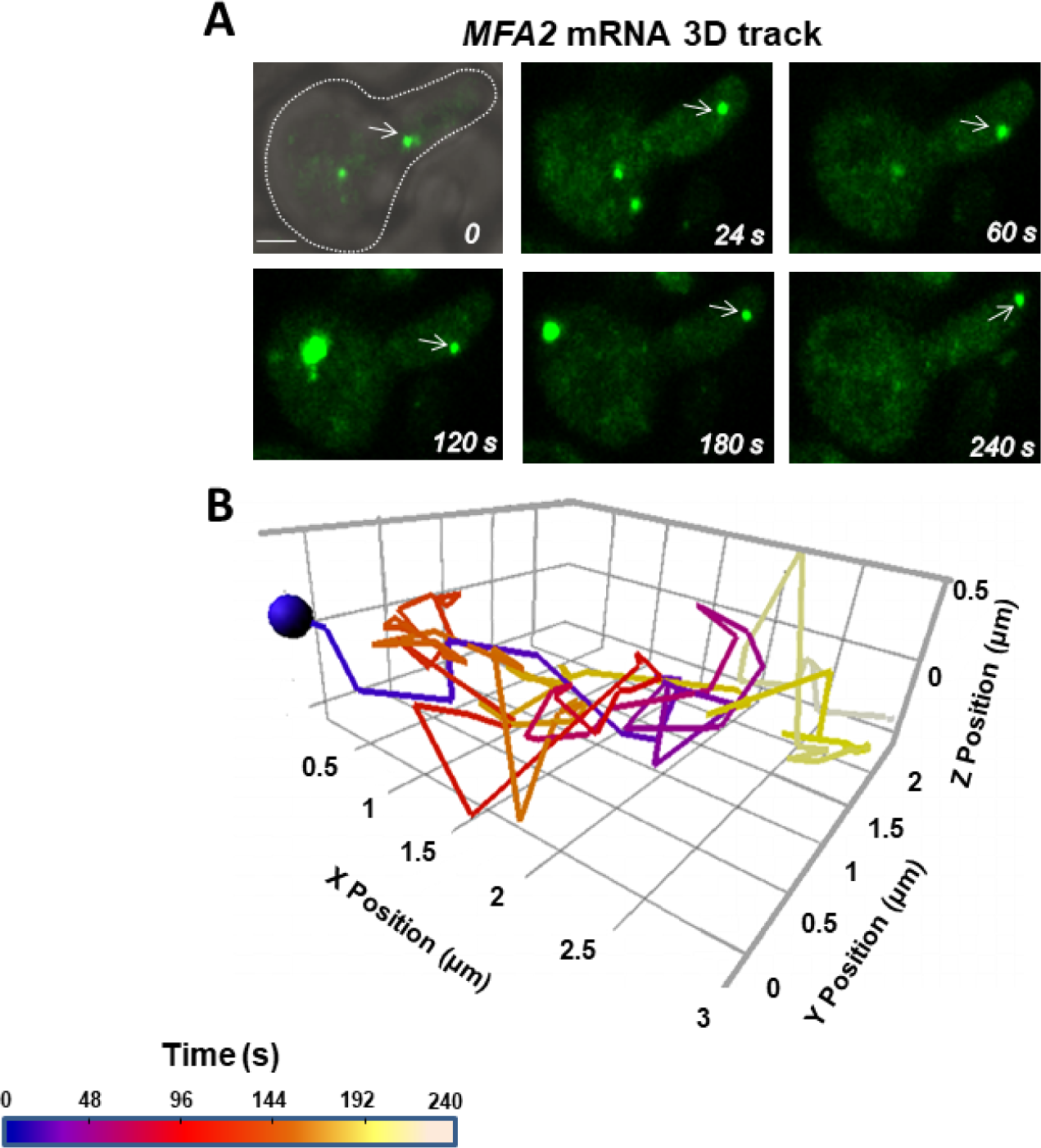
A translocating *MFA2* granule made numerous twists and turns in the z-direction. **(A)** Cells expressing labeled *MFA2* mRNA (green) and treated with α-factor for 2 hours were recorded in 3D by confocal microscopy. White arrows indicate the motile, low-intensity, transport mRNA granule analyzed in (B). Scale bar = 2 µm. **(B)** Although route mapping in 3D was coarse grain compared to that in 2D, frequent detours in the z-direction were apparent. The color scale refers to the time. Track displacement rate was 0.01 μm/s, mean velocity was 0.09 μm/s.

In both cases (2D and 3D), the translocational movement did not take the shortest route from the mother to the shmoo tip. The granule typically went through many twisting and rotary motions close to its initial location before performing one or more large directed jumps toward the shmoo (S1_Movie.mov, S3_Movie.mov, and S4_ Movie.mov). However, it became clear that “straight” segments in 2D usually involved deviations in the z-direction.

Interestingly, granules arriving at the shmoo tip did not lose the ability to move; they instead reverted to local oscillation similar to what was typical prior to a jump (e.g., Fig. 3). Normally, pheromone binding to G-protein coupled receptors stimulates polarized growth in one direction, towards the source of the pheromone gradient established by the nearest cell of opposite haplotype [38]. Granules did not stop moving at their destination in this experimental setting, possibly because the α-factor was present in the medium all around the cell. In addition, directed granule movement may have had to wait for further maturation of the shmoo, since shmoo extension was incomplete in many of the cells under study.

### Physical movement parameters of *MFA2* and *PGK1* granules

Next, we tested the randomness of granule movement by examining the nature of motion on a fine time scale, supported by known physical parameters. Two-dimensional analysis was more appropriate, because greater time resolution was required. Fluorescent labeled *MFA2* and *PGK1* mRNAs were imaged in yeast that were treated with α-factor for 2 hours to induce shmoo growth. The cells were recorded at 14 -18 frames/s for 60 s. Figure 5A-E shows representative *MFA2* and *PGK1* granule tracks of the various movement types.

**Fig. 5.**
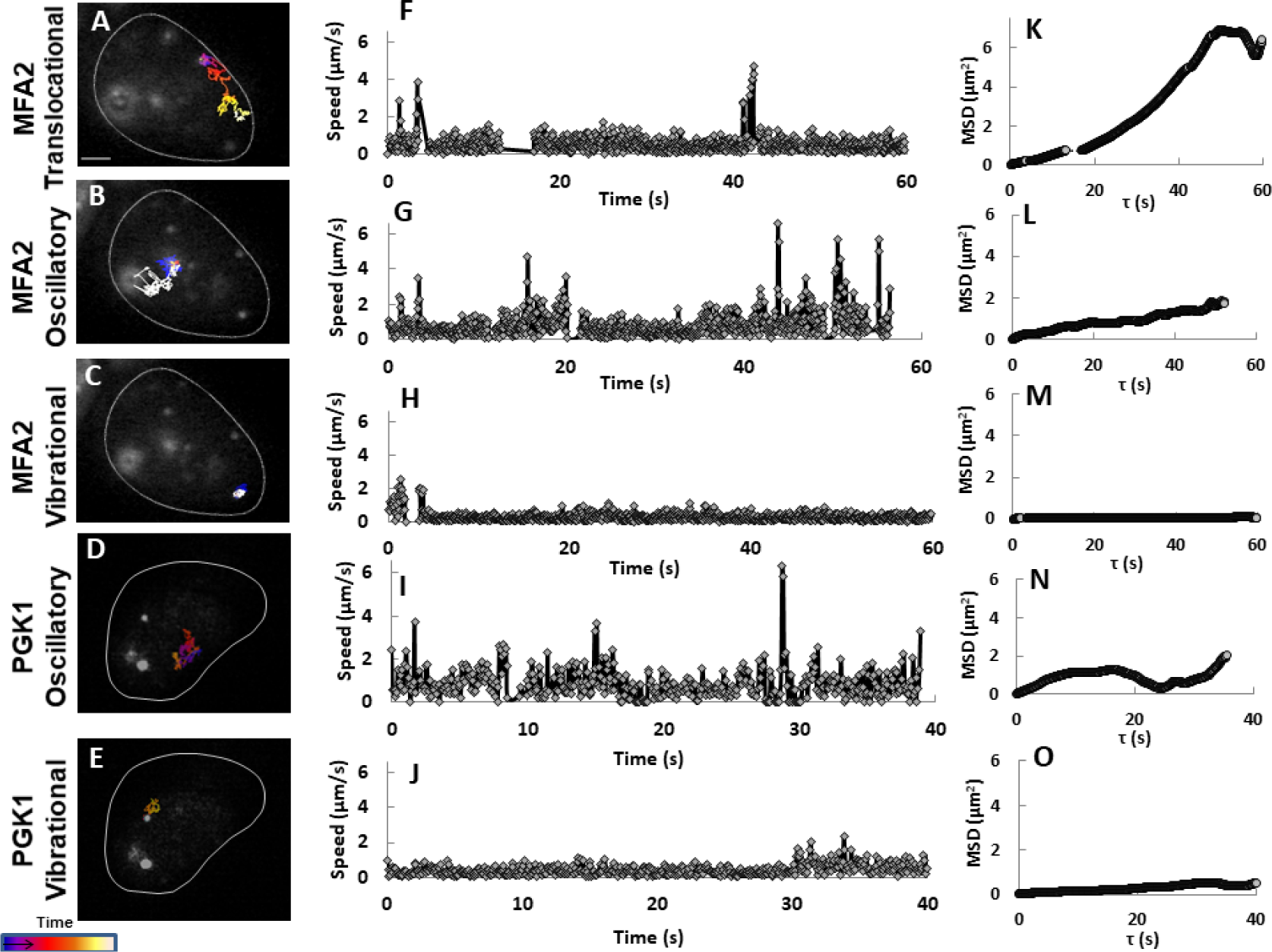
MSD rather that velocity distinguished the different types of movement. Three *MFA2* **(A-C)** and 2 *PGK1* **(D-E)** granules were tracked in yeast cells treated with pheromone for 2 h before movies were taken in 2D by fluorescence microscopy. The interval between frames was 0.055-0.07 s. Scale bar = 1.5 µm applies to (A-E). **(F-J)** Speed at a given time point was given as the average of the local velocities determined for 3 neighboring time points, where local velocity was the calculated as the distance traveled over two frames divided by one time intervals. **(K-O)** Mean Square Displacement (MSD) was diagnostic for random and nonrandom translocation as well as corralled movement.

Two additional parameters were analyzed: local velocity and mean square displacement (MSD). Local velocity at a given time point was estimated from the 2D position immediately preceding and the position immediately following that time point. Both vibrating *MFA2* and *PGK1* granules showed start-stop movement and relatively slow local velocities when moving, with a few bursts up to 2 μm/s (Fig. 5H, J). Interestingly, there were high-intensity *MFA2* mRNPs (size 0.5 µm) containing 40-50 mRNA copies, located in the shmoo (Fig 5C, H and M). These granules were Mating Bodies that play a role in controlling the translation of aF. Based on MSD analysis the Mating Bodies were attached within the shmoo tip. Oscillatory RNPs granules of both mRNA types had similar local velocities fluctuating between 0 and 2 μm/s, that were punctuated by high-speed jumps of up to 5-6 μm/s (Fig. 5G, I). We found that small *PGK1* granules did not translocate to the shmoo, but *MFA2* granules undergoing translocation behaved like oscillating granules in terms of local velocity (Fig. 5F), although they sometimes reached speeds as high as 8 μm/s. However, the MSD profiles associated with these two types of movement were completely different.

The MSD represents an ensemble average in the sense that it measures the spreading of many particles, characterized by the spatial average of squared particle position [39], (see Methods). According to Brownian motion theory, MSD is proportional to the time during which the movement is random. If movement is directed, then the MSD function tends to follow a parabolic or an exponential function over time. When the MSD profile decreases it is a “corralled” motion due to some local potential [40]. Figure 5K shows that while an *MFA2* granule was undergoing translocation (Fig. 5A), the MSD function was parabolic. In contrast, one oscillatory granule (Fig. 5B) showed a linear MSD graph (Fig. 5L) that indicated random movement. Hence, the MSD parameter demonstrated that despite the similar distributions of local velocities between the two types of granules, the nature the movement was quite different: in figure 5A it was directed transport until it reached the shmoo and in figure 5B, it was random walk. The *PGK1* mRNA granule in figure 5D exhibited oscillatory movement within a confined area hence the MSD profile remained below a maximal value and included a segment with a negative slope. Vibrational *MFA2* and *PGK1* granules (Fig. 5C, E) did not change their locations by more than 0.9 μm and were marked by MSD profiles with very low slopes (Fig. 5M, O). Thus, vibrational granules were tethered or corralled and probably not engaged in very slow, directed movement.

### Variable movement of *MFA2* granules within a single track

*MFA2* granules undergoing translocation often appeared to engage in a different type of movement initially (e.g., Figs. 3, 5A). To explore these granules further, we examined their changes in position relative to the starting point in the x-and y-directions separately over time, the angle of movement, and a ψ parameter, that indicated the probability that a granule would stay within a circle of maximum displacement. (See Methods, S1_Appendix.pdf). Figure 6 shows the analysis of the *MFA2* granule in figure 5A as it moved from the cell body to the shmoo, focusing on two sections of the track. The first consisted of 100 frames in the initial, 5.5s segment of the track (green square in Figure 6A) in which the granule oscillated within a limited region. The second section (blue square in Fig. 6A) includes 100 frames in which the *MFA2* mRNA granule translocated. These two tracks, made by the same granule, differed substantially with regard to all of the parameters analyzed. Possibly, this RNP was subjected to a switch in regulation due to an exchange of its RBP components.

The green section was marked by a lack of order in terms of positional change (Fig. 6B) and angle of movement (Fig. 6C). Vantage visualization of the granule route revealed a track displacement of only 0.21 µm (Fig. 6E) and a linear MSD profile (Fig. 6F). Furthermore, the parameter log(ψ) exceeded the threshold (see Methods), suggesting a high probability of the granule remaining within a region of maximum displacement of 0.11 µm, in this track fragment (see Methods) (Fig. 6D). Hence, this small, low-intensity *MFA2* granule initially seemed to move randomly without venturing far from its starting point.

**Fig. 6.**
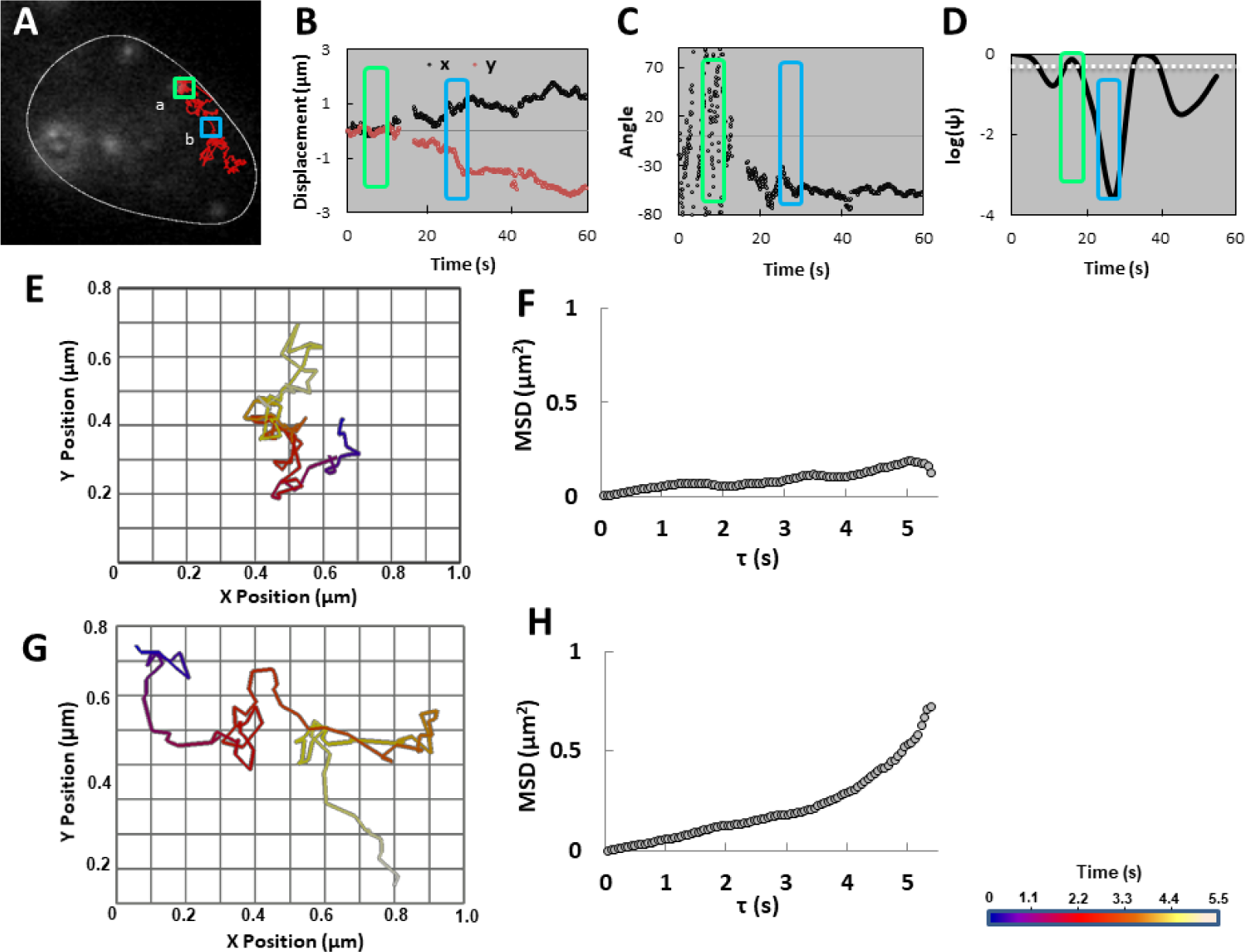
Transitions in type of movement were observed in the same track. **(A)** Two sections of the track for the granule in figure 5A, marked by the green and blue squares, were analyzed further in 2D in panels (B)-(F). **(B)** Granule displacements on the x- and y-axes as a function of time were minimal at first (green box), but then increased in magnitude over time (blue box). **(C)** Angle of the movement relative to its position at the start of the observation period as a function of time was initially random (green box), then turned several times (e.g. blue box), before settling on a particular directionality. **(D)** Log(ψ) of the track was calculated in segments of 100 frames as a function of the time. The white dashed line represents the Log(ψ) threshold=-0.3. Log(ψ) falling below the threshold was indicative of directed motion. **(E), (G)** Vantage visualizations are shown for the track segments marked by the green and blue boxes in (A), respectively. The color scale refers to different time periods lasting 5.5 s in these two panels only. **(F), (H)** MSD parameter was calculated for the track segments marked by the green and blue boxes in (A), respectively.

In contrast, in the blue section there was a fairly steady increase in displacement away from the starting point in both x- and y-axes (Fig. 6B) and a constant rate of change in the angle of movement (Fig. 6C) overall. Finer time resolution revealed greater complexity with some backward jumps along one axis and a few very brief epochs that mimicked vibrational movement (Fig. 6G). The log(ψ) parameter was much lower than in the green section predicting a lower probability of the granule staying within a circle of maximum displacement of 0.14 µm, in this track fragment (Fig. 6D). The highlight was the MSD plot for this section of movement with a parabolic profile indicating directed rather than random motion. It is noteworthy that even when analyzed over the entire 60 s period during which movement was nonuniform and riddled with local wandering (producing a variable log(ψ), Fig. 6D), jitters and jumps with velocities up to 4 µm/s (Fig. 5F), the MSD profile was still parabolic for most of the track (Fig. 5K).

These findings belie movement that was much more complex than was apparent from lower time resolution imaging. Different factors regulate the mRNA movement in different parts of the cell and probably there were numerous underlying mechanisms. Since the actin cytoskeleton is required for mRNA localization [10] and actin cables grow along the cell [41], we can assume that when the granule was moving in short steps, it was not carried by a motor protein on the actin [39], but in parts of the route where there were larger jumps, some motor protein may have participated in the process and supported the higher velocity of movement.

### *MFA2* granules co-transported with PB proteins to the shmoo tip

Since *MFA2* mRNA co-localizes with PB proteins in the shmoo tip in α-factor treated cells [27], the question arose as to whether the mRNA trafficked as PBs, or whether they formed PBs once they arrived at the destination. To find out, the *MFA2* mRNA labeling system was used along with an additional plasmid for Dcp2p-RFP, a PB component coupled to a different fluorophore. In three independent experiments, 100 granules in 20 cells were analyzed separately for movement of *MFA2* mRNA (GFP fluorescence) and PBs (RFP fluorescence). A cell with seven granules is shown in figure 7 and in S6_ Movie.mov. The bulk of granules in the center and 4 separate granules (3 white arrows and 1 red arrow) co-labeled for PB protein and *MFA2* mRNA. Co-labeled granules displayed all three types of motility and the mean track velocities determined for each granule by following GFP and RFP separately, showed good correspondence: vibrational (0.05 µm/s), oscillatory (0.15 µm/s) and translocational (0.13 µm/s), indicating that PB proteins accompanied *MFA2* mRNA in the transport, localization, storage and sorting processes (Table 3). Figure 7B shows the route of one *MFA2* mRNA-PB as it translocated from cell body to shmoo. Moreover, this track was very reminiscent of the translocational tracks examined in figures 1 and 3, after taking into account the different rates of frame capture.

**Table 3.** Velocities of *MFA2* granules with and without PB protein

| Type of motility | *MFA2* mRNA with Dcp2p | Dcp2p | *MFA2* mRNA |
| --- | --- | --- | --- |
| Vibrational | 0.05±0.04 | 0.05±0.04 | --- |
| Oscillatory | 0.15±0.01 | 0.15±0.03 | 0.22±0.07 |
| Translocational | 0.13±0.03 | 0.13±0.03 | 0.23±0.01 |

Mean velocity = track length/track duration, in units of $\mu\text{m/s}$ . 85 granules, 0.2 to 0.8 $\mu\text{m}$ in diameter, were analyzed in 16 cells. PB granules were defined by the presence of Dcp2p (see text). Data are given as mean ± SEM.

**Fig. 7.**
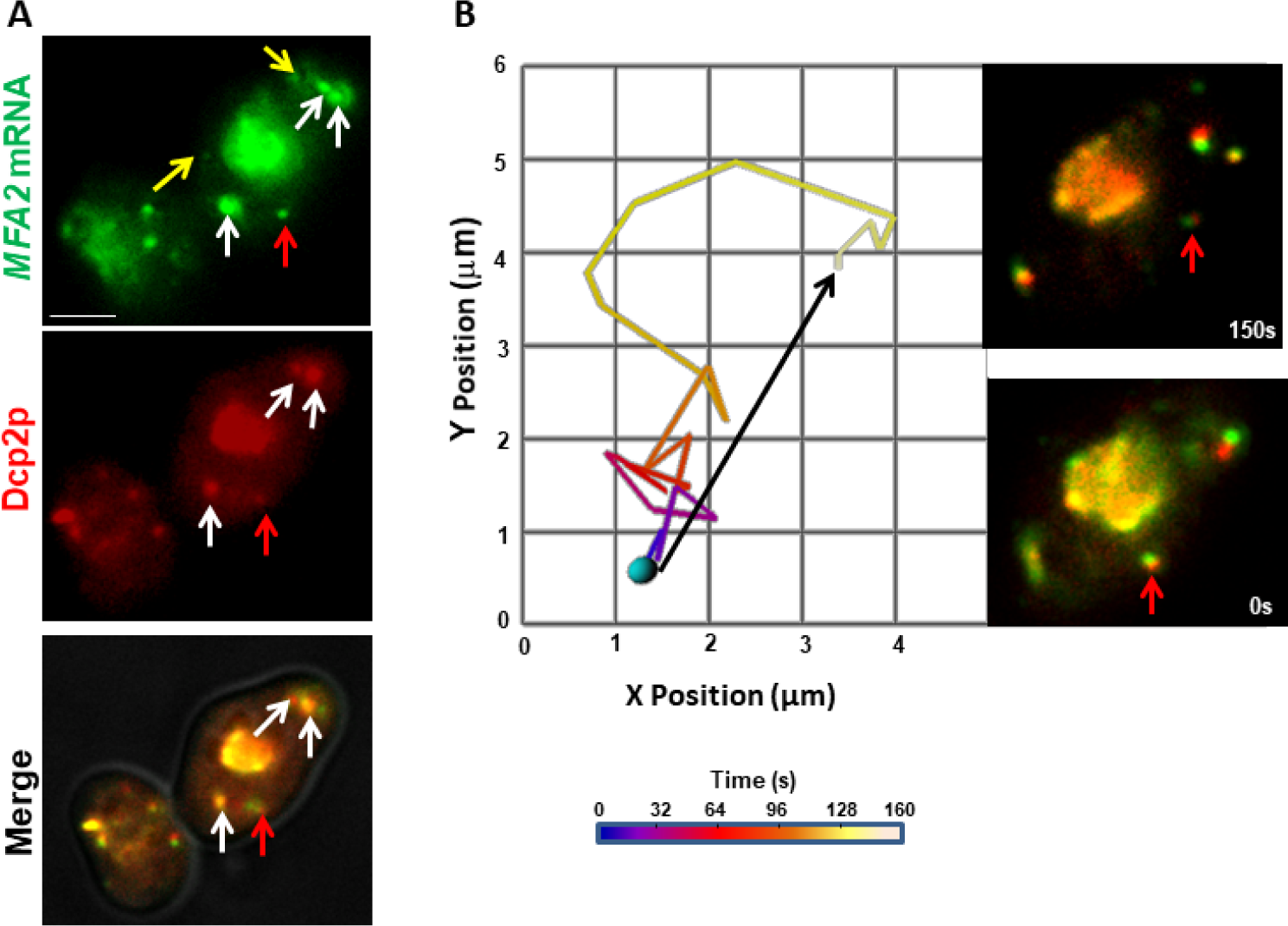
*MFA2* mRNA and the PB marker co-transported to the shmoo tip. **(A)** *MFA2* mRNA (green) co-localized with Dcp2p-RFP (red), a PB marker, in many granules. Several small co-labeled granules are indicated by the red and white arrows, while granules with *MFA2* alone are marked by yellow arrows. Scale bar = 5 µm. **(B)** The 2D track is shown for the granule in (A) indicated by the red arrow. The track displacement of this translocational granule (black arrow) over an observation period spanning 150 s, was 3.8 µm. Inset: the two markers remained co-localized during the entire track.

Some *MFA2* granules were not co-labeled by the PB marker (Fig. 7A, yellow arrows). Because Dcp2p is a core PB component [42], we considered these granules not to be PBs. The composition of such granules should be determined in a future study. *MFA2* mRNA granules lacking Dcp2p showed comparable oscillatory movement but a translocational movement with a two-fold faster mean velocity (Table 3), compared to granules of the same size that bound Dcp2p. Interestingly, vibrating mRNA granules missing Dcp2 were not observed. Granules with Dcp2p alone were not observed indicating that all PBs contained *MFA2* mRNA. Therefore, we can conclude that translocation of *MFA2* granules could occur without PB formation, but that PBs participated in at least a fraction of *MFA2* mRNA transport to the shmoo tip (See Discussion). It follows that PB formation affected the modes of movement available to a granule.

### Myo4p involvement in *MFA2* mRNA transport

Deletion of Myo4p, the main motor protein in budding yeast, did not prevent *MFA2* mRNA localization or inherence of cortical ER to the shmoo (S1 Fig. panel B). Here we investigated whether the absence of Myo4p changed the movement of *MFA2* granules in a discernible way. *MFA2* mRNA was labeled in a yeast strain lacking Myo4p (Δmyo4). After cells were treated with α-factor for 2 hours to induce shmoo formation, 2D tracking analysis was performed on images recorded at 14-18 frames/s for 60 s. Although the localization percentage of *MFA2* granules at the shmoo was similar to that in WT yeast, granules translocating from cell body to shmoo were never observed in mutant cells (Fig. 8 and S7_ Movie.mov). All movements were either vibrational (Fig. 8D, G), oscillatory in which the granules moved in a confined area (Fig. 8B, E), or oscillatory with larger displacements not directed toward the shmoo (Fig. 8C,F). According to the displacement parameter the latter can be considered as translocational, but the overall track MSD profile was linear (See also Discussion). The movement types were verified by the analysis of MSD. The diffusion coefficient was less than or equal to 1 µm^2^/s. Similar to WT, “corralled motion” indicated by a negative slope of the MSD profile was observed (Fig. 8C, F). Thus, while *MFA2* mRNA localization did not strictly require the presence of Myo4 motor protein, the mRNA movement was affected. It implied that one mode of transport to the shmoo was blocked by Myo4p deletion, so the *MFA2* mRNA that did localize to the shmoo: translocated as very small granules not detected by our methods, were possibly carried by a different motor protein, or used another mechanism entirely.

**Fig. 8.**
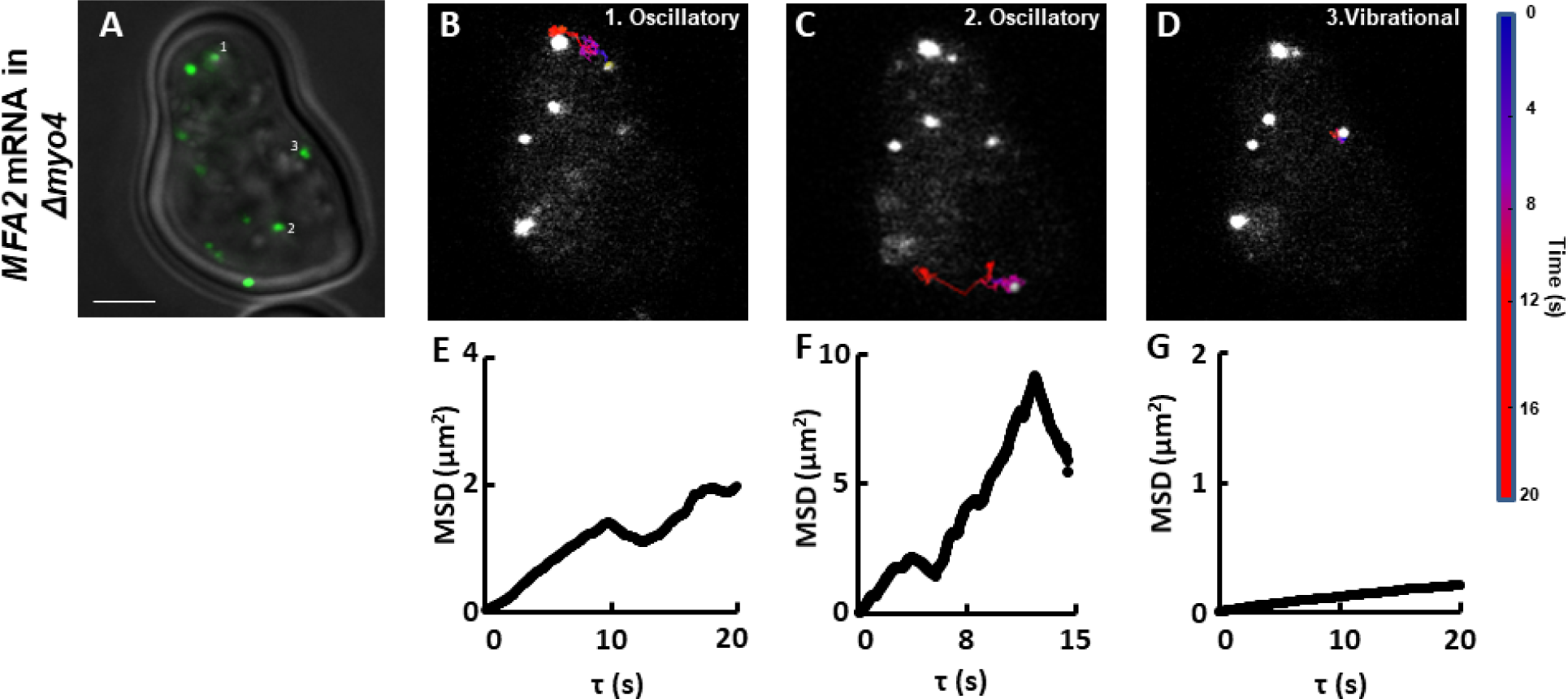
*MFA2* granules in Δmyo4 yeast did not exhibit fast translocational movement to the shmoo. **(A)***ΔMyo4* cells expressing labeled *MFA2* mRNA were treated with pheromone for 2 h and recorded at 0.055-0.07s time intervals for 60 s. Scale bar = 2 µm also applies to (B-D). **(B-D)** Tracks of 3 particles in mutant cells were limited to two motility types: vibrational and oscillatory. The color scale refers to time. (E-G) Each MSD profile corresponds to the track presented above it in panels (B)-(D).

Table 4 compares the granule velocities of the WT strain versus the mutant *Δmyo4* strain. Granules undergoing vibrational and oscillatory movements showed very similar velocities between strains. This finding emphasizes that these two types of granule movements were unaffected by the absence of the Myo4p motor, and only the ability to move in a directed fashion to the shmoo was lacking. Yet somehow, *MFA2* granules still localized in the shmoo.

**Table 4.** Motility types of *MFA2* granules in *Δmyo4* vs WT strains

| Type of motility | Velocity in WT ($\mu\text{m/s}$) | Velocity in $\Delta Myo4$ ($\mu\text{m/s}$) |
| --- | --- | --- |
| Vibrational | 0.08±0.07 | 0.08±0.06 |
| Oscillatory | 1.0±0.2 | 1.1±0.3 |
| Oscillatory with large displacement | 1.7±0.3 | 1.6±0.4 |
| Translocational | 1.5±0.4 | ---- |

Mean velocity= track length/track duration, in units of $\mu\text{m/s}$ . 100 granules, 0.2-0.8 $\mu\text{m}$ in diameter, were analyzed in 20 cells. Data are given as mean ± SEM.

**Table 5.** *S*. *cerevisiae* strains used in the study

| Strains | Genotype | Source |
| --- | --- | --- |
| ySA061 | *MATa, ura3, his3, leu2* [pPS2037, pRP1187] | Aronov et al., 2015 |
| ySA056 | *MATa, ura3, his3, leu2* [pSA03, pRP1187] | This study |
| ySA022 | *MATa, ura3, his3, leu2* [pSA03, pRP1187, pRP1152] | This study |
| ySA118 | *MATa, ura3, his3, leu2, myo4: NEO*; [pSA03, pRP1187] | This study |

## Discussion

Here, for the first time, we examined *MFA2* and *PGK1* RNP transport in high resolution and characterized their motile properties in aF treated cells.

### Comparison of 2D and 3D tracking

Research into the mRNA dynamics in yeast has been conducted mainly in two dimensions. Notably, there are two 3D studies: the first simultaneously monitored in space two types of mRNAs, *ASH1* and *IST2* [19], and the other one performed ARG3 mRNA tracking using a double-helix point spread function and included in-depth physical analysis of the different types of movement [6]. We now report three-dimensional tracking of mRNA movement in yeast undergoing profound structural changes associated with mating. In the present study, we compared *MFA2* mRNA surface tracking (2D) to spatial tracking (3D).

Our 3D tracking confirmed the occurrence of the three types of *MFA2* granule movement: vibrational, oscillatory and translocational (Fig. 1), which were previously described in 2D tracks [27]. Only two types of movement: vibrational and oscillatory, were observed for the *PGK1* granules that distribute throughout a yeast cell (Fig. 2). 3D tracking required collecting information from the entire cell volume and thus was necessarily performed at a relatively slow rate of image capture, 2-4 s/frame. The average velocity of translocating *MFA2* granules in 3D was estimated to be 0.15 µm/s (Fig. 4) and was lower as compared to 2D analysis. This estimate is two times lower than the velocities determined on different mRNAs in budding yeast in 3D in studies for which the temporal resolution was greater [6,19]. The question then arose as to whether our estimate of 3D velocity reflected reality.

Currently, much greater temporal resolution is possible with 2D imaging, 0.55-0.07 s/frame, and sensitivity is greater, so small, ultra-fast granules of low fluorescence intensity can be tracked [27]. The average velocity of these translocating granules in 2D was 1.5 µm/s (Table 2), consistent with previous reports [24,27]. However, track displacement rates in 2D and 3D, which provide very coarse grain estimates of velocity, were similar (Table 2). Therefore, the disparity between the average velocities of granules was mainly attributed to the difference in time resolution of the methods. Consistent with this interpretation, velocities for several granules tracked in 2D were recalculated for positions determined at 2 s intervals, and the resultant “intermediate” coarse grain values converged with the slower velocities determined for other cells by 3D tracking. Nevertheless, 3D tracking did reveal a substantial amount of movement in the z-direction, indicating that the rapid jumps observed in 2D tracking were not always as straight as they appeared. It remains to be seen whether faster velocities will be measured with even greater time resolution and since our 2D values ignored movement in the z-direction, the translocating mRNA granule velocity determined from our 2D approach must be considered to be a lower limit.

### Complexity of *MFA2* mRNA transport

*MFA2* and *PGK1* mRNPs fluorescent labeling using a U1A plasmid-based system has shown differences not only in number and sizes of the granules for both RNP types, but also distinct dynamic behavior including motility properties (trajectory displacement, and length, Figure 1-2, Table 1). However, we do not exclude the possibility that these differences were dependent on transcript length; *MFA2* is short (300 bp), whereas *PGK1* is long (1500 bp). Therefore, the type(s) of RBPs incorporated in the mRNP complexes were probably distinct and may have conferred different properties.

The MSD parameter distinguishes intentional and directed movement from random movement [43]. A completely random walk (Brownian motion) is characterized by a zigzag trajectory of variable track displacement and a near-linear MSD profile with a shallow slope. In our study, directed movement resulted in a parabolic MSD profile. Using this and other physical parameters in the present study, we demonstrated that *MFA2* mRNA translocational movement from the cell body to shmoo was extremely circuitous. Furthermore, transitions between directed transport and diffusive motion could be discerned. Statistical tests of the switching within a single trajectory between directed and confined modes of motion reveals that 21% are confined-restricted stationary movement, 4% are directed, and the remaining 75% are classifiable by default as consistent with diffusion [6]. In budding yeast, *ASH1* mRNAs reaching the bud neck suddenly jump directly to the bud tip [19]. We identified in mating yeast, many translocating granules undergoing a change in the type of movement (e.g., Figs. 3, 6; S3_Movie.mov, S5_ Movie.mov).

Changes in speed may occur after a change in granule composition. Sometimes we observed small granules coalescing to create larger complexes (see S3_ Movie.mov, big central granule starting at the 10th second). Previously, actin patch assembly was observed in yeast that had formed a shmoo [41] and actin networks participate in mRNA localization as the shmoo grows [10]. Slight movement of an mRNA granule may occur due to oscillation of the actin fiber or result from tiny steps along an actin fiber that has some curvature [44]. In addition, assembly and disassembly of actin patches are very quick during the establishment and maintenance of cell polarity, with a patch lifetime of 11±4 s [41]. Hence, granules might also pause at actin branching points, sites of actin remodeling, or in places where there are actin discontinuities. Furthermore, we never observed two granules following the same route to the shmoo, consistent with volatility in the “cell’s roads” and the existence of a wide network of roads.

The directed movement of *ASH1* mRNA from the mother cell to the daughter cell uses the Myo4p motor protein [7]. The velocity of transit can be affected by the number of motor molecules that transfer the cargo [28]. Although *MFA2* mRNA localized in the shmoo, we failed to see mRNA granules being transported to shmoo in yeast lacking Myo4p. We could offer several explanations. First, as we previously mentioned, translocation is a relatively rare phenomenon, so in the absence of Myo4p, we just did not observe it. This possibility seems improbable given that we did observe translocation in WT yeast. Second, similar to the cortical ER inheritance mechanism that depends on the cytoskeleton but also relies on anchorage [45], it is possible that some mRNA moved through the cytoskeleton without motor proteins and localized at the shmoo due to a strong affinity of the complex to that specific location. Transport and anchoring are distinct processes, for example, directed mRNA movement occurs from the cell body to bud and then granules switch to confined movement within the daughter cell [19]. Third, *MFA2* mRNA may have transported as individual transcripts, with fluorescence too weak for us to resolve [37], and aggregated at the shmoo as a visible granule. Finally, we see hints of a very fast transit mechanism involving sudden, big leaps of oscillatory with large displacement granules.

### Which motor protein participated in *MFA2* mRNA transport?

Class V myosins have biochemical and structural properties suitable for actin-based transport. In yeast, five genes of the myosin family are known to be expressed. Two are class V myosin; Myo2 and Myo4 are motor proteins that move along actin cables within the cell [46].

Myo4p protein participates in the transport of mRNA and ER in budding yeast [29,47] as well as the localization of *SRO7* mRNA in mating yeast [14]. Furthermore, PB transport is dependent on the Myo4p/She2p RNA transport machinery in budding yeast [23]. We demonstrated that localization of large *MFA2* granules at the shmoo as well as cER inheritance persists without Myo4p (S1 Fig.). In the present study, granules moved at the same velocities in mutant yeast lacking Myo4p. *MFA2* mRNA did accumulate at the shmoo, however, directed movement of the granules from the cell body to the shmoo was never observed (Fig. 8). Myo4p was identified as a principal motor of *ASH1* mRNA transport in budding yeast operating with a velocity of ∼0.3 μm/s [7]. At first sight this rate falls short of the measured *MFA2* mRNA granule velocity of ∼1.5 µm/s for granules that undergo translocation in mating yeast, but those former calculations relied on videos acquired at 1s intervals, which were 18 times slower that our acquisition rate in 2D. Therefore, Myo4p could mediate the fastest *MFA2* mRNA transport, but did not exist as the exclusive means of transport. Future study is required to figure out the alternative mechanisms of *MFA2* mRNA transport.

Myo2p is more typical for transporting vascular organelles, such as Golgi or peroxisomes, with velocities of movement of 0.15-0.45 µm/s [46]. It also mediates rapid, exocytotic-vesicle transport into the bud with a velocity of 3 µm/s [48]. Fungal RNA could be transported by vesicle in the case of extracellular vesicle-mediated export of nucleic acids [49], but *MFA2* mRNA granules are considered to be cytosolic bodies [27] that translate and undergo a series of biogenesis steps occurring in the cytosol [32,50]. Thus, we rule out the possibility that *MFA2* mRNA was packed into vesicles and then transported by Myo2p. Myo2p can associate with a large, RNP particle(s) and form a PB that is distinct from actively translating polysomes and exosomes [51], but confirmation that Myo2p provides a means for directed transport of mRNA will require additional study.

### PB involvement in *MFA2* mRNA transport

In mammalian cells, PB movement is not usually direct and is mediated by microtubules [52]. In budding yeast, the movement of PBs is undirected, dependent on Myo4p/She2p RNA transport machinery, and occurs along the actin cytoskeleton from the mother to the daughter cell. In addition, we showed 100% co-localization between Dcp2p and high-intensity granules of *MFA2* mRNA and 44% co-localization between low-intensity granules. We recorded the common transport of the PB component, Dcp2p, together with *MFA2* mRNA in yeast treated with pheromone α (Fig. 7). The observation shows that in oscillatory and translocational modes, granules moved along with and without the PB protein, whereas in vibrational mode, granules all co-localized with Dcp2p [53]. The finding suggested two scenarios that were not mutually exclusive. First, a fraction of fast *MFA2* granules translocated to the shmoo independent of PBs, whereas another pool moved as a PB complex. Second, an initial stage of the RNP complex formation excluded Dcp2p protein, but it joined later, just prior to movement toward and retention at the shmoo. During its life cycle PB size is not constant; the PB can undergo several cycles of growth and decomposition [54]. Hence, during its movement in the cytoplasm, RNP complex composition is variable and different factors may control its movement at different stages of the journey.

Based on our observations, we propose the following model of *MFA2* mRNA fate in shmoo forming cells (Fig. 9). Cytoplasmic mRNAs are regulated at the post-transcriptional level and transcript localization is one important means for translation control. There are three types of cytoplasmic *MFA2* mRNPs that determine cytoplasmic fate (vibrational, oscillatory and translocational), as well as a fourth RNP type called the *MFA2* MB located in the shmoo tip that is responsible for aF translation and subsequent secretion of aF from the cells. The *MFA2* RNPs are distinct in size, subcellular location, and numbers of RBPs associated with them, and therefore, they have different motility properties. First, immotile RNPs, located in the cell body, manifest as vibrational mRNA granules. From their properties and location, it seems likely that they sit on and attach to peri-nuclear ER. The second type of the RNPs move quickly, although their movement is restricted to the cell body sub-region, therefore they comprise the oscillatory mRNP type. These RNPs are partially attached to some structure in the cell body, we assume that these type of granules can be attached also to tubular ER, which are characterized as dynamic structures that change their size and location during shmoo growth and are pulled to the bud/shmoo tip of the cell. [55,56]. *MFA2* RNP velocity can be dependent on the growth of tubular ER to the shmoo. Interestingly, that tubular ER appears to exclude ribosomes, which means that translationally inhibited RNPs or PBs can be associated with it [55].

**Figure 9.**
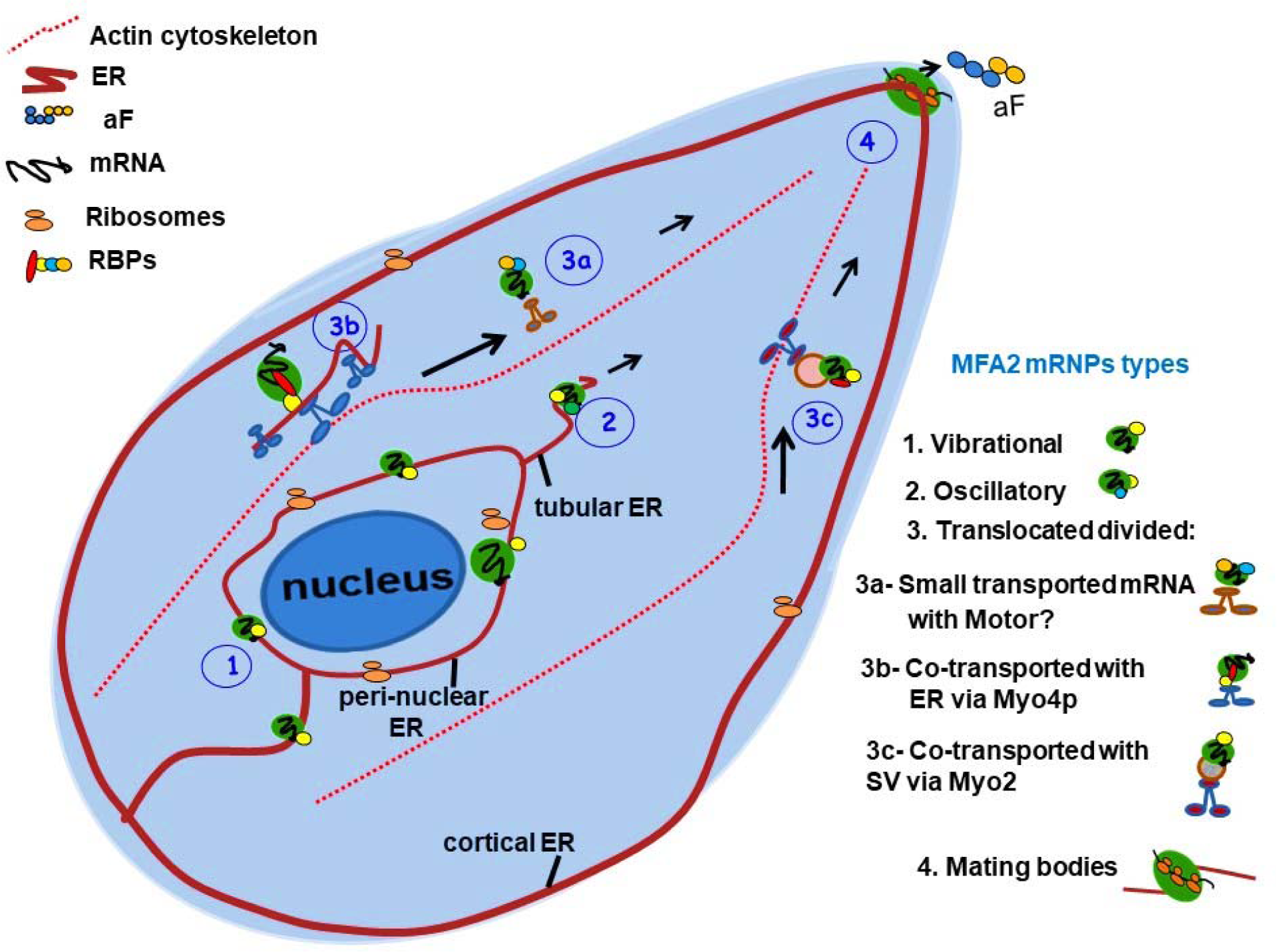
Model of *MFA2* mRNA fate in alfa pheromone treated cells

The third type of *MFA2* RNP is the translocated granule that moves to the shmoo. According to our results, some *MFA2* mRNA granules transported to the shmoo via Myo4p though actin and showed complex movement. Since RNP delivery to the shmoo occurs in the absence of Myo4 motor, we propose that transport can be accompanied by tubular ER pooled to shmoo, that then branches out to form new contacts in multiple directions with the PM to reestablish [45]. Therefore, tubular ER can be an alternative means for *MFA2* mRNA transport to the shmoo.

We also do not exclude the possibility that translocated mRNPs can be delivered to shmoo together with perinuclear ER derived secretory vesicles SV. These vesicles accumulate early during shmoo growth [57]. These hypotheses should be investigated in future research.

## Materials and methods

### Yeast strains and growth conditions

The *S*. *cerevisiae* strains and plasmids used in this study are listed in Tables 5 and 6. Cells were grown at 25–30°C in synthetic complete medium to the early exponential growth phase. For selection, the appropriate amino/nuclear acid was omitted from the media, since its expression was directed by the exogenous plasmid (see Table 6, yeast marker).

**Table 6.** Plasmids used in the study

| Plasmid<sup>a</sup> | Genotype | Yeast | Source |
| --- | --- | --- | --- |
| pRP1187<sup>b</sup> | *GPDp-U1A-GFP* | LEU2 | P. Silver<sup>c</sup>; Brodsky and Silver, 2002 |
| pMC313 | *GPDp-MFA2-U1Ax16-3'UTR of MFA2* | URA3 | Aronov et al., 2015 |
| pPS2037 | *PGK1p-PGK1-U1Ax16-3'UTR of PGK1* | URA3 | S. Aronov<sup>d</sup>; Aronov et al., 2015 |
| pSA03 | *MFA2p-MFA2-U1Ax16-3'UTR of* | URA3 | This study |
| pRP1152 | *DCP2p-DCP2-RFP* | HIS3 | R. Parker<sup>e</sup> |

<sup>a</sup>All plasmids, unless otherwise indicated, contained CEN and therefore were present in one or two copies per cell.
<sup>b</sup>Multicopy 2μ plasmid.

### Labeling system/plasmids

For in vivo mRNA labeling, two plasmids were transfected into yeast cells. One plasmid contained the chimeric U1A-GFP protein. The second plasmid included the target gene with a UA1-binding site (loops). Centromeric plasmid pSA03 contained an *MFA2* promoter and its open reading frame, with a 3⍰-UTR that included 16 repeats of the UA1-binding site for in vivo fluorescent labeling of its mRNA molecules in living cells [58]. A full length *MFA2* 3’UTR fragment was amplified by PCR and inserted between SacI and Not1 site digest of pRP1193 using: forward primer, 5’-cggactagtccgccgcggcTTTTTGACGACAACCAAGAG where capital letters are the *MFA2* 3’UTR and small letters denote a SpeI restriction site and reverse primer, 5’-atttgcggccgcggcGCGGAGGGAAAGGCGTATC-3’, where capital letters are the *MFA2* 3’UTR and small letters denote a Not1 restriction site. The sequence was verified by Sanger sequencing. Centromeric plasmids carrying *PGK1*, whose 3⍰-UTR also contained 16 repeats of U1A-binding site (pPS2037), pU1A-GFP (pRP1187) and DCP2p-RFP (pRP1152), were kindly provided by Prof Roy Parker (Howard Hughes Medical Institute, University of Colorado Boulder, Boulder, CO, USA) (Table 6).

### α-factor treatment

Stock solution of α-factor (Sigma-Aldrich) consisted of 1 mg/ml in methanol. Cells were grown in synthetic complete medium to early exponential phase and treated with 3 nM α-factor for 2 h. In some cases, the culture was collected by centrifugation and resuspended in a six-fold smaller volume just before the addition of α-factor. Only fresh cell samples of low concentration were inspected under the microscope (<15 min) to avoid adverse effects (e.g., starvation, hypoxia) of incubating them in between the slides.

### Image processing and analysis of time series movies

The 3D and long time-lapse videos of mRNA granules and proteins in mating yeast were taken at room temperature (∼25°C) by confocal microscope using a 63x/1.40 NA objective (LSM 700; Carl Zeiss). In these experiments, cells were typically imaged in 3D (six - seven z-planes per time point), at 0.5 µm steps. The time point duration 2-4 sec 2D recording with two fluorescence filters was done with a fluorescence microscope (Olympus IX81, Objective: 100 x PlanApo, 1.42 numerical aperture). The following wavelengths were used: for GFP, excitation at 480 nm and emission at 530 nm; for RFP, excitation at 545 nm and emission at 560–580 nm. There was a 2.5s delay between the green and red fluorescence measurements. For imaging two fluorophores in a cell, sequential screening was performed to avoid overlapping with the LSM image. The AxioVision LE64, Zeiss software was used for producing 3D movies. Labeled *MFA2* and *PGK1* mRNA were tracked and the movies presented as a z-series compilation of 7- 10 photographs in a stack (0.5 µm) using the AxioVision LE version 4.8.2.0, Zeiss software. The quantitative co-transport analysis of the *MFA2* mRNA/Dcp2p-RFP over a period of time was carried out in Imaris 7.4.2 software (Bitplane AG).

The short videos of *MFA2* and *PGK1* granules were produced at room temperature (∼25°C), using the inverted motorized fluorescent microscope with a 100x/1.30 objective (Axio Observer.Z1; Carl Zeiss), an LED light source (Colibri) and a high speed camera (Zeiss HS;Carl Zeiss). Videos were made using AxioVision Rel. 4.8 imaging program (Carl Zeiss). The camera had a peak QE of more than 90%, making it a near-perfect detector that could capture images at high speed with an unequalled signal-to-noise. This high sensitivity allowed us to follow the low intensity, faster, transported granules in the cell.

For presentation of the movies, the 3D image sequences were transformed into a time sequence using the maximum projection option to 3D. Videos were processed using Imaris 7.4.2 software (Bitplane AG). Statistical data analysis was performed using Excel 2010 version 14 (Microsoft). The final images were prepared with Photoshop CS6 version 13.0.06 (Adobe).

### Tracking analysis of *MFA2* or *PGK1* granules

The 2D and 3D trajectories of *MFA2* and *PGK1* granules that were acquired using a confocal microscope (Fast Acquisition, LSM 700; Carl Zeiss) at room temperature (∼25°C) were built using the semi-manual mode of the Imaris 7.4.2 program. The granules chosen for analysis had at least 40 visible time points without unviable gaps exceeding 3 consecutive time points. 3D movies had several z-stacks in which only one or two of them caught the granule. For analysis, we superimposed all z-stacks at each time point, then drew a straight line between time points. This meant that there were epochs when the granule movement could not be determined accurately and the track was assumed to be smooth, but was in reality, most likely to be kinked.

Local or instantaneous velocity was calculated using an interval of 3 time points. The displacement, duration, track length, and types of tracking were analyzed according to intensity and position of the mRNA granules in the cell using the same program. Mean velocity was calculated as the total track length divided by the observation period. Except where specifically stated otherwise, track displacement and track length were given for a standardized track duration of 60 s. In cases for which track durations greater than 60 s were available, values were determined for 60±5s segments and then averaged. Vantage images show the track pathway in 2D/3D scale using Imaris 7.4.2 program (Bitplane).

MSD, log(ψ), and angle of *MFA2* and *PGK1* granular motion were analyzed based on x, y, z and time parameters, obtained from track building. MSD profiles were calculated for each trajectory by a classical Brownian motion algorithm [40], and plotted as a function of τ. τ is the lag time between the two positions taken by the particle. For log(ψ) analysis, the route was divided into sub-tracks of 100-time frames (t_w_)_;_ in each sub-track the maximum displacement value was defined (R). The diffusion coefficient (D) was calculated from the derivative of the MSD function. Then probability, log(ψ) was calculated by taking the linear approximation of Dt_w_/R^2^ which will be valid for R < sqrt(Dt).

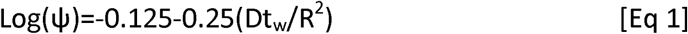

The measured probability was compared to a set threshold (ψ = 0.5). A probability ψ > 0.5 would indicate an increased particle distance within a finite time, having lost the ability to be confined [59]. More details on the fitting algorithm can be found in Text SI.

## Acknowledgments

We thank Susan Michaelis for the generous gifting of yeast strains, and Clint L. Makino for his invaluable comments on the manuscript.

## Supporting Information

**S1 Fig.**
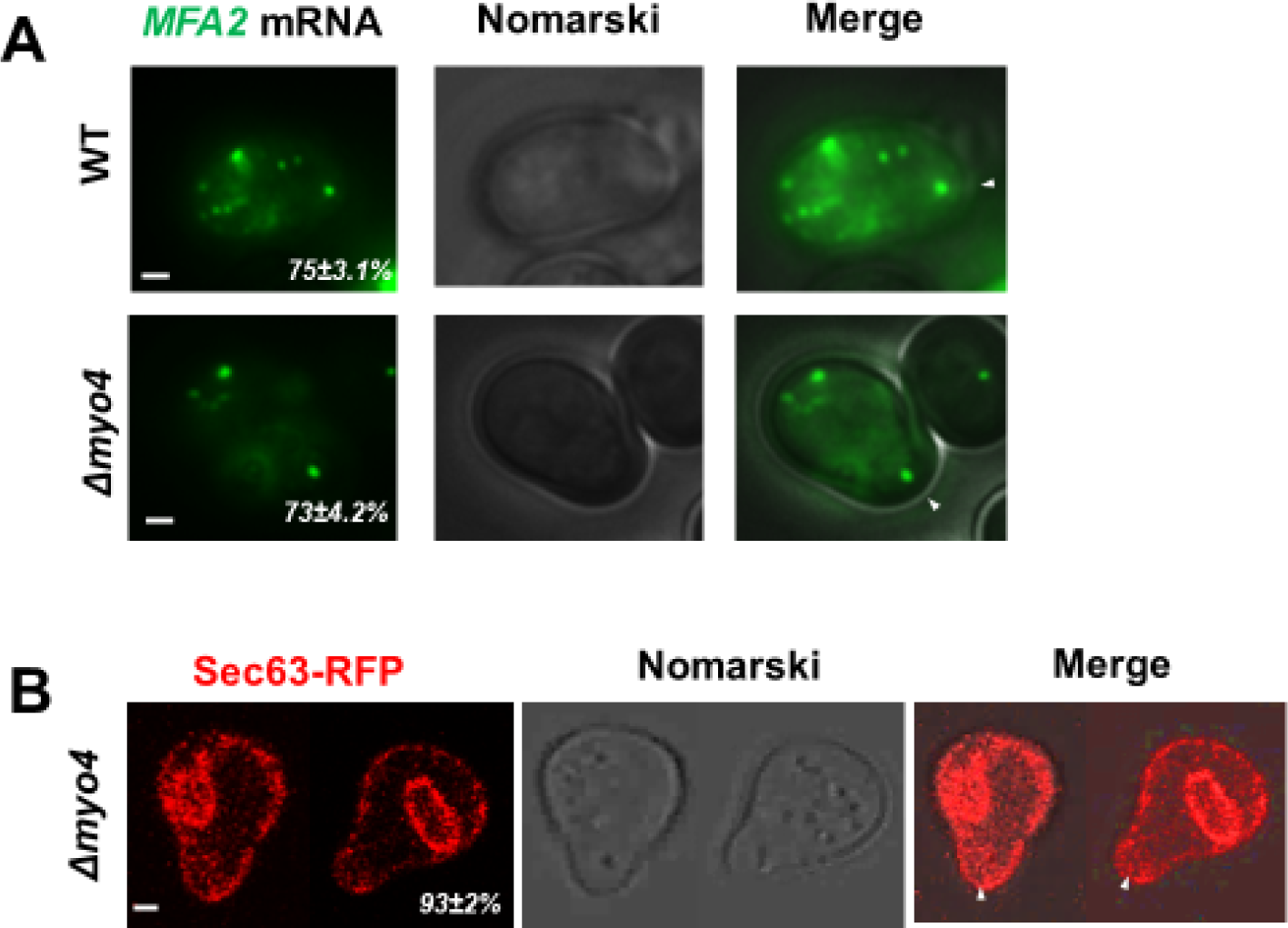
*MFA2* mRNA transport and cortical ER inheritance to shmoo is not dependent on Myo4p motor protein. WT and *Δmyo4* strains grown until the logarithmic phase were treated with α-pheromone for 2 h. (A) Example of *MFA2* mRNA distribution within the cell. The distribution of *MFA2* mRNA was analyzed by fluorescence microscopy by performing 6 z-stacks with a depth of 0.5 µm for each cell. White arrows indicated *MFA2* mRNA localization in the shmoo tip. Bar, 1 µm. (B) *Δmyo4* yeast cells expressing ER marker, Sec63-RFP, analyzed by confocal fluorescence microscopy by performing 6 z-stacks with a depth of 0.5 µm for each cell. Cortical ER inheritance was tested in 90 cells in three independent experiments. White arrows indicated cER co-localized with plasma membrane throughout the cell and specifically in the shmoo tip. Bar, 1 µm.

**Movie 1S**. *MFA2* **granules showed diversity in movement using 3D tracking analysis**. Images captured in 3D of a cell co-expressing *MFA2*-U1A and U1A-GFP (ySA056) following treatment with 3 nM α-factor for 2 h. Images were taken every 4 s (entire z-stack) for 97.9 s and replayed as a high-speed movie. Elapsed time is tracked in the bottom left corner. The track of one granule is highlighted with colored line segments as it moved to the shmoo. The color sequence corresponds to the time tracked on the color bar on the bottom right. Figure 1 shows representative snapshots from this movie. The highlighted granule in the movie is particle 3 in figure 1.

**Movie 2S** *PGK1* **granules showed vibrational and oscillatory, but not translocational movement using 3D tracking**. Cells co-expressing *PGK1*-U1A and U1A-GFP (ySA056) were treated with 3 nM α-factor for 2 h. The tracks of three granules are highlighted with colored line segments. Images were captured every 3.5 s (entire z-stack) for 97.6 s. The movie shows a fast-forward version, at a rate of 5 frames/s. Elapsed time is shown on the bottom left and the timing of the granule tracks are followed on the color bar on the bottom right. Figure 2 shows representative snapshots from this movie.

**Movie 3S Two-dimensional time-lapse analysis captured the rapid movements of the low intensity** *MFA2* **granules**. Cells co-expressing *MFA2*-U1A and U1A-GFP (ySA056) were treated with 3 nM α-factor for 2 h. Images were captured at a rate of 0.055 s/frame using fluorescence microscopy and presented in slow motion, at a rate of 10 frames/s. The elapsed time is shown on the bottom left and is tracked on the color bar on the bottom right. Figure 3 shows representative snapshots from this movie.

**Movie 4S. Three-dimensional tracking of an** *MFA2* **granule shows its circuitous route from the cell body to the shmoo tip**. Cells co-expressing *MFA2*-U1A and U1A-GFP (ySA056) were treated with 3 nM α-factor for 2 h and followed with time-lapsed, confocal microscopy. Images were captured every 2 s (entire z-stack) for 243 s, and shown as a movie at 2 frames/s. The track of a translocating granule in the central cell that had a well-developed shmoo was charted with colored line segments. The elapsed time is shown on the bottom left and is tracked on the color bar on the bottom right. Figure 4 shows representative snapshots of the central cell from this series of images.

**Movie 5S. Subtypes of the** *MFA2* mRNA **transport were observed in the same track**. Cells co-expressing *MFA2*-U1A and U1A-GFP (ySA056) were treated with 3 nM α-factor for 2 h. Images in 2D were captured using fluorescence microscopy every 0.055 s during a 1 min period. The movie shows a fast-forward version, at a rate of 50 frames/s. The elapsed time is shown on the bottom left and is tracked on the color bar on the bottom right. Figures 5 and 6 show representative snapshots of the cell in the middle of the field, from this movie.

**Movie 6S. *MFA2* granules co-transported with Dcp2p-RFP (PB marker) in α-factor activated cells**. Cells co-expressing *MFA2*-U1A, U1A-GFP and the P-body marker, Dcp2p-RFP (ySA22), were treated with 3 nM α-factor for 2 h prior to time-lapse photography with images captured every 5 s for 155 s, using the fluorescence microscope and played back at 2 frames/s. The elapsed time is shown on the bottom left and is tracked on the color bar on the bottom right. Figure 7A shows representative snapshots from this movie. The track of the granule indicated with the red arrow in figure 7A is followed with colored line segments.

**Movie 7S**. *MFA2* **granules in** *Δmyo4* **yeast cells did not appear to translocate to the shmoo**. Mutant *myo4Δ* cells co-expressing *MFA2*-U1A and U1A-GFP (ySA056) were treated with 3 nM α-factor for 2 h. Images were captured using the fluorescent microscope with 0.055 s/frame during 28.8 s. The movie is shown at a rate of 20 frames/s. The elapsed time is shown on the bottom left and is tracked on the color bar on the bottom right. Figure 8A-D shows representative snapshots from this movie.

## S1 Appendix

### Diffusion coefficient and diffusion equation for the probability of finding the particle within a defined area

The diffusion equation in cylindrical coordinates is:

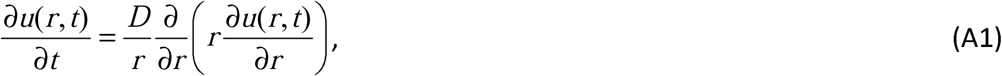

where *u*(*r,t*) is some dynamical variable. In our case, it may be a concentration (density) of particles at *r* and at a moment *t*. The coefficient *D* describes the diffusion. Obviously, at a point *r* at a moment *t*, only one particle can be located. We may say ‘concentration’ in the sense that there are a number of particles in the vicinity of *r* in a volume small compared with the characteristic volume of the problem, and this number can be divided by this small volume.

If we consider some circular area (0,*R*) bounded by a wall then the boundary condition for our problem is *u*(*R,t*) = 0. This means that the particles which initially were in the above-mentioned circle are confined (“corralled”) within this circle. With some arbitrary initial condition *u*(*r*,0) = ⍰(*r*) and using the method of variable separation, *u*(*r,t*) = *F*(*t*)*G*(*r*), we obtain the solution:

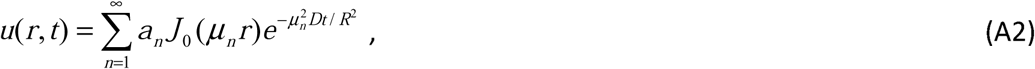

where ***J*_0_** is the zeroth order Bessel function, **μ_*a*_** is its n th root, and the condition of the solution at *r* = 0 is used. The unknown coefficients should be found from the initial and boundary conditions. With the new variable *x* = *r*/*R* the initial condition reads

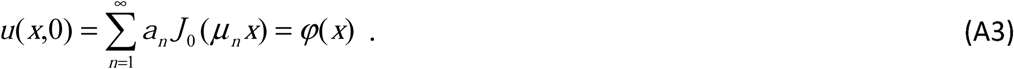

Multiplying eq.(A3) by ***xJ*_0_**(**μ*m***) and integrating over the region [0,1], with the use of well-known identities we obtain

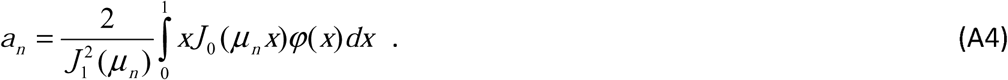

If *u* has the sense of a concentration and we are interested in the dynamics of a single particle (granule) in the area bounded by ***x* ≤ 1** during a sufficiently long time, then the concentration is 1/*πR*^2^ or 1/*π* in normalized form. Therefore, ⍰(*x*) = 1/*π*. Thus, we finally obtain

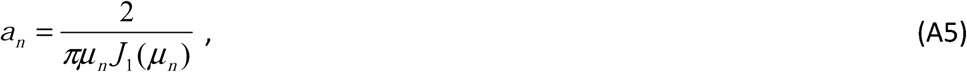

Where *J*_1_ is the first order Bessel function. The solution now reads

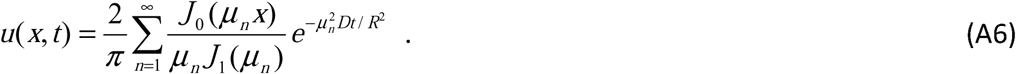

The probability of finding the particle in the area bounded by [0,*x*] is

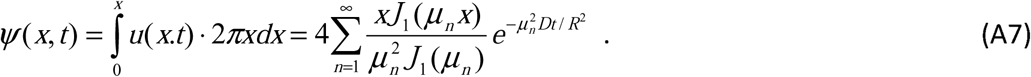

Truncation of these series by retaining the first five terms gives the accuracy better than 2%. According to Einstein-Smoluchowski’s theory, the mean square distance (MSD) that a particle diffuses from its initial position by Brownian motion will be **⟨ x**^2^**⟩ − *Dt*** For experimental trajectories of proteins in biological cells, the calculated MSD as a function of time deviated from linearity, indicating non-Brownian motion [60]. Such motion can be caused by factors that drag out or “corral” the particle.

We have normalized the particle position *r* by the radius *R*. If we consider this radius as the expected scale of the “corral” where the particle is confined, then a reasonable size of this area may be a tenth of the cell radius, i.e. *R* ∼ 1-2 µm (the maximum length of the cell with developed shmoo was 12-15 µm). Hence, what remains to be estimated is the diffusion coefficient *D*. Following Einstein’s formula which is based on the Stokes’ law of viscous drag force, the diffusion coefficient can be approximated as

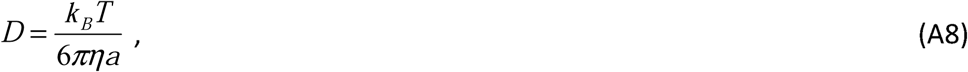

where *k*B is Boltzmann’s constant, *T* is absolute temperature, *η* is the dynamic viscosity coefficient of the medium, *a* is the diameter of the particle (thought of as spherical). Using the viscosity coefficient in water *η* ∼ 0.8 mPa·s and *R* ∼ 0.1 µm, an upper limit of D/*R*^2^ ∼ 3.5×10^−3^ s^−1^ was obtained.

#### Calculating the granular angle of motion

The angular position of *MFA2* mRNA were calculated by the equation:

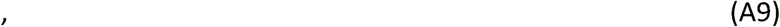

Where α in the Δx and Δy were the changes in position from the beginning of the track.

